# Normalization of Single-cell RNA-seq Data Using Partial Least Squares with Adaptive Fuzzy Weight

**DOI:** 10.1101/2024.08.18.608507

**Authors:** Vikas Singh, Nikhil Kirtipal, Songwon Lim, Sunjae Lee

## Abstract

Normalization of single-cell RNA-seq (scRNA-seq) is a crucial step in downstream analysis, where raw data are adjusted to correct unwanted factors that prevent the direct comparison of expression measures. scRNA-seq data exhibits a multivariate relationship between transcript-specific expression and sequencing depth that a single scale factor cannot address. A partial least squares (PLS) regression was performed to accommodate the variability of gene expression in each condition, and upper and lower quantiles with adaptive fuzzy weights were utilized to correct unwanted biases in scRNA-seq data. The present approach was compared using real and simulated datasets across various state-of-the-art performance measures.

## I. Introduction

Effective pre-processing and normalization play a vital role in studying gene behavior under distinct biological conditions in both scRNA-seq data [1]–[3] and high-throughput sequencing data [4]–[6]. Gene behavior under these conditions can vary due to unwanted biases caused by transcript length, GC content, dropout, capture efficiency, sequencing depth, and other biological or technical factors [7]. The implicit normalization approach aids downstream analysis, such as mitigating inflated false positives in differential expression analysis, by correcting of sequencing depth, cell-to-cell differences in capture efficiency, and other technical factors [8].

In the literature, various methods have been presented for within-sample and between-sample normalization of bulk RNA-seq data [9]. The most commonly used methods, which have been highly successful, are the trimmed mean of M values (TMM) [10], DESeq [11], and median ratio normalization [12]. However, these methods do not perform well for scRNAseq data due to the prevalence of zero-expression counts. The zero-expression counts make scRNA-seq data extremely sparse, resulting a zero geometric mean (GM) in DESeq and undefined M values in TMM. Additionally, the unbalanced library sizes of abundant and rare cell populations in scRNAseq may violate the non-DE hypothesis, leading to bias [13].

In recent years, various methods have been proposed for scRNA-seq normalization, falling into two broad classes: global scaling factors and gene-specific scaling factors. The most commonly used approaches for scRNA-seq include multiplying by a constant factor [1], BASiCS [14], SAMstrt The code and experimented data are available on the GitHub platform: https://github.com/vikkyak/bIbW/tree/main [15], the gamma regression model (GRM) [16], pooling normalization [8], robust normalization of scRNA-seq data (SCnorm) [17], and PsiNorm, which models scRNA-seq data using a Pareto distribution [18]. A significant bias with the global scaling factor is that it accommodates the transcript-specific expression and sequencing depth relationship by using a common scale factor for all genes in a cell, leading to over-correction when this relationship is rare across genes. However, gene-specific scaling factors can also introduce bias during normalization, particularly when uniform count-depth relationships are assumed across cells, which can vary substantially when marker genes are highly expressed in one cell type but not in others. Recently, kernel-weighted-average robust normalization for single-cell RNA-seq data (scKWARN) [13] was developed to correct known or unknown technical factors without relying on explicit count-depth relationships or data distributions. It generates a pseudo-expression count for each cell using fuzzy technical neighbors through kernel smoothing.

The present approach overcomes biases due to library size, dropout, RNA composition, and other technical factors and is motivated by two different methods: pooling normalization [8], which pools cells with similar library sizes, and scKWARN [13], which does not rely on specific count-depth relationships or data distributions. In this approach, the scRNAseq data is clustered, and PLS analysis is performed on each cluster or condition. To preserve data variability, correlation is calculated for each condition, and each condition is multiplied by the average correlation to minimize differences in library size before normalization. Each condition is then divided into upper and lower quantiles using the mid-quantile function of Qtools, and a confidence interval is obtained using the confint.midquantile function. PLS analysis is conducted using the pls package in R to obtain the first component from the upper and lower quantiles that exhibits highest variance. This first component is used to measure the adaptive fuzzy weight of the upper and lower quantiles to determine the expression level of the genes, which is then used to calculate the scale factor and normalize the data, as described in Figure 1.

**Fig. 1:**
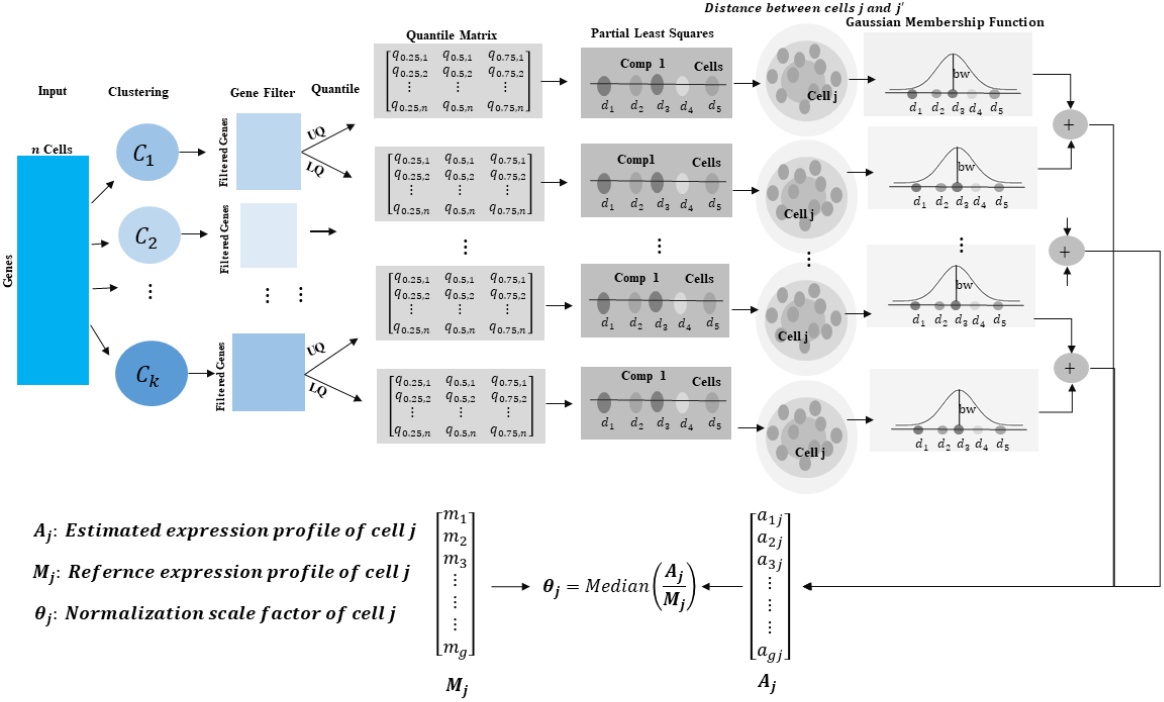
Overview of sample normalization of scRNA-seq data using partial least square with adaptive Gaussian fuzzy weight.

The rest of the paper is organized as follows: Methods of the present approach is discussed in Section II. Results on real and simulated data is discussed in Section III. Computational performance in IV, Finally, Section V concludes the paper.

## II. Material AND Methods

The present approach first utilizes unsupervised clustering to pool the cells. Then, without relying on specific count-depth relationships or data distributions, it generates a pseudo-expression count for each cell. This approach is divided into five distinct steps as follows:

1. In the first step, if the conditions of datasets are not available. The present approach, cluster the datasets using <u>quickCluster</u> function of <u>scran</u> library in R to group the dataset into multiple clusters. However, any clustering algorithm can be utilized to group the datasets into a random number of groups to proceed to the next step.
2. The number of variables (genes) in each group may differ, and the nature of the variables (quantitative or qualitative) can vary from one group to another, but the variables should be of the same type within a given group [19]. To address this issue, we used Partial Least Squares (PLS) regression [20], which simultaneously considers multiple sets of variables and balances the influence of each set. We ran <u>pls</u> with one component and calculated the cross-correlations between the components associated with each condition. To balance each group or condition, we multiplied it by the average cross-correlation.
3. In the third step, technical similarity for each condition is estimated using the first PLS component with 25^*th*^, 50^*th*^, and 75^*th*^ percentiles of the upper and lower quantiles of non-zero expression count in cells, using the <u>midquantile</u> function from the <u>Qtools</u> package in R. To elaborate, assume an scRNAseq data has *m* genes and *n* cells. Let *y*_*gj*_ be the expression count of *g* gene in the *j* cell for *g* = 1, …, *m*, and *j* = 1, …, *n*. Define *P*_*j*_ = *{g* : *y*_*gj*_ *>* 0, *g* = 1, …, *m}* is the subset of genes with non-zero expression counts in cell *j*. Let *Ql*_*j*_ = [*q*_0.25,*j*_, *q*_0.50,*j*_, *q*_0.75,*j*_] and *Qu*_*j*_ = [*q*_0.25,*j*_, *q*_0.50,*j*_, *q*_0.75,*j*_] represent the upper and lower log nonzero expression counts, respectively, with each being a 3-D vector containing 25^*th*^, 50^*th*^, and 75^*th*^ percentiles of non-zero expression counts with log-transformed in cell *j*. Let *dl* = [*d*_1_, *d*_2_, …, *d*_*n*_] and *du* = [*d*_1_, *d*_2_, …, *d*_*n*_] are the first PLS component of *Ql* and *Qu*, respectively.
4. In the fourth step *dl* and *du* are utilized to measure the technical variability among cells by estimating the weight 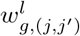 and 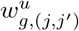 as shown in Figure 1, step, Gaussian membership function. The lower and upper weights are defined as follows:

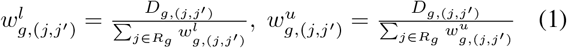

Where

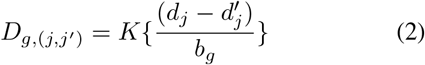

K(.) is the Gaussian kernel function, and *b*_*g*_ is bandwidth eastimated by a kernel density estimate in R package KernSmooth. The pseudo expression count for cell *j* are written as

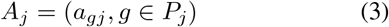

Where

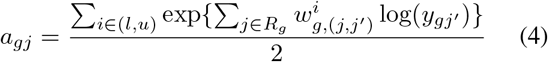

5: The final step creates a reference expression count for each cell *j*, denoted as *M*_*j*_:

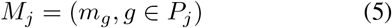

Where

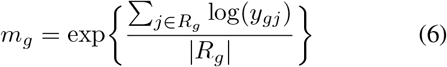

Here, *m*_*g*_ is the geometric mean (GM) of gene *g* across the subset of cells *R*_*g*_ where the gene is expressed. In other words, for each non-zero count gene *g* in cell *j*, the process involves determining the subset of cells express this gene (*R*_*g*_) and finding the GM of the gene expression across *R*_*g*_. It is important to note that *M*_*j*_ is calculated based on non-zero count of genes in the cell *j*, making *M*_*j*_ specific to cell *j*. The pseudo count of each cell is then compared to its specific reference count to find the size factor of each cell. The present approach estimates the normalization factor using a median ratio, as follows:

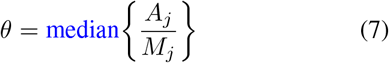

## III. Results AND Discussion

The performance of the proposed (Pro) approach is assessed in terms of F1 score, log-fold change, root mean square error (RMSE), and the proportion of differentially expressed genes, and is compared with other state-of-the-art methods, including Relative Counts (RC), scran, sctransform, scKWARN, and PsiNorm, using three real scRNA-seq datasets and two simulated datasets described below.

### A. Real Datasets

#### 1) PBMC33K

The first dataset used in this study is PBMC33K, generated by 10x Genomics, which comprises 33,148 human peripheral blood mononuclear cells and 32,738 genes. Similar to scKWARN, we randomly extracted 1,000 B cells and generated two cell populations for a comparable analysis. A fixed percentage of genes was randomly selected and scaled with distinct factors to increase the proportion of DE genes in these two cell populations. The present approach was tested on moderate and strong DE genes with balanced and unbalanced cell populations. In the case of moderate DE, the percentage of genes varied from 5% to 15% in each cell population; in strong DE, the percentage of genes varied from 15% to 30%. The balanced case had 1,000 cells in each population, while the unbalanced case had one population with 1,000 cells and the other with 400 cells. Model-based analysis of single-cell transcriptomics (<u>MAST</u>) was utilized to identify the DE genes [21]. In all cases, the present approach was compared with scKWARN, RC, sctransform, scran, and PsiNorm using performance metrics such as the F1 score and RMSE with true fold change. For moderate DE with balanced and unbalanced settings, as shown in Figure 2, the present approach achieved comparable or better performance in F1 scores and RMSE values. Figure 2 also shows that the present approach accurately estimates the true log-fold change while varying the percentage of DE genes within the cell population. For strong DE with balanced and unbalanced settings, as shown in Figure 3, the present approach achieved performance comparable to scKWARN and outperformed the other normalization methods in F1 scores and RMSE values.

**Fig. 2:**
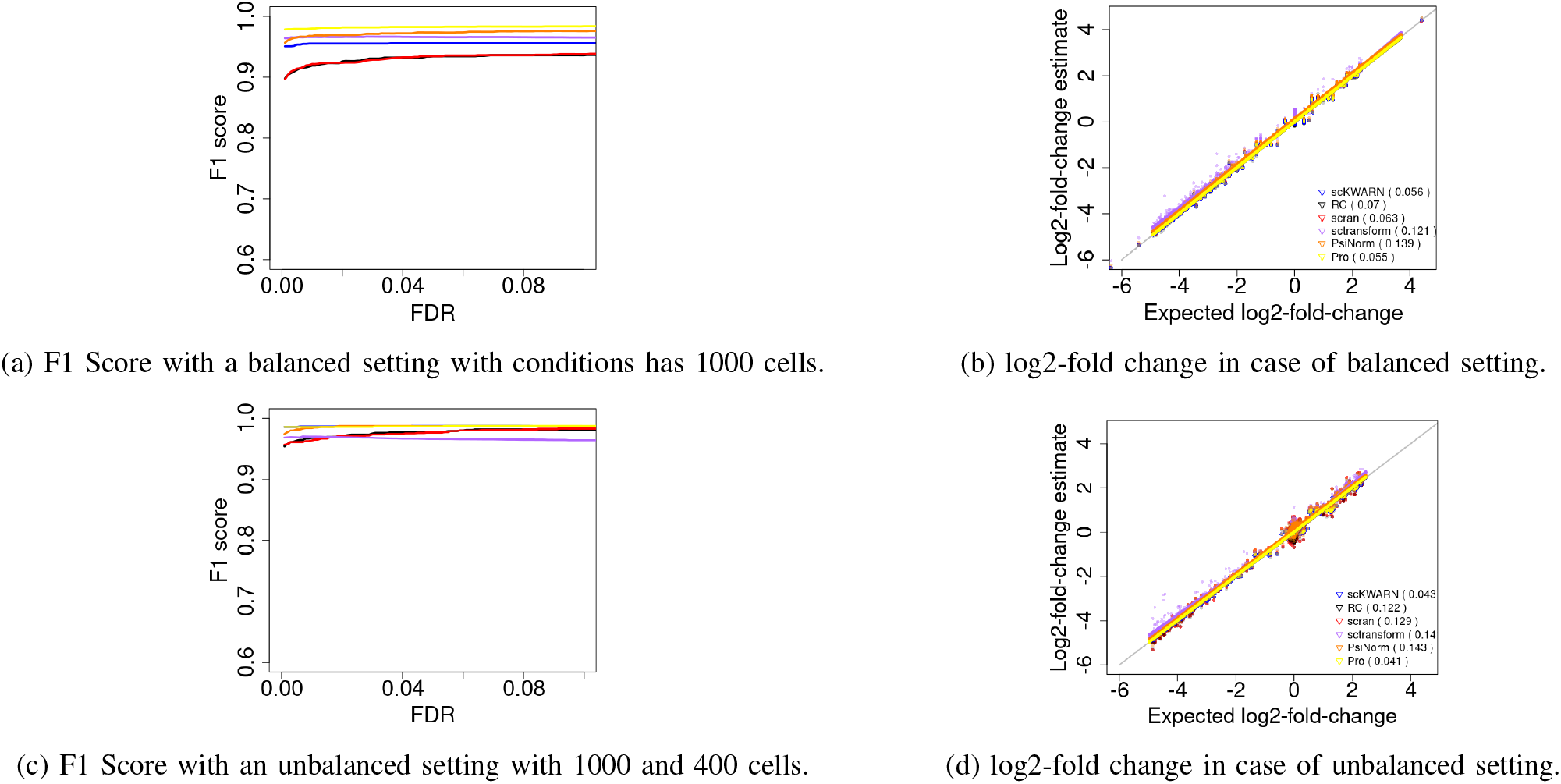
Performance analysis with state-of-the-art on PBMC33K dataset with Moderate (10%) Differentially Expressed Genes.

**Fig. 3:**
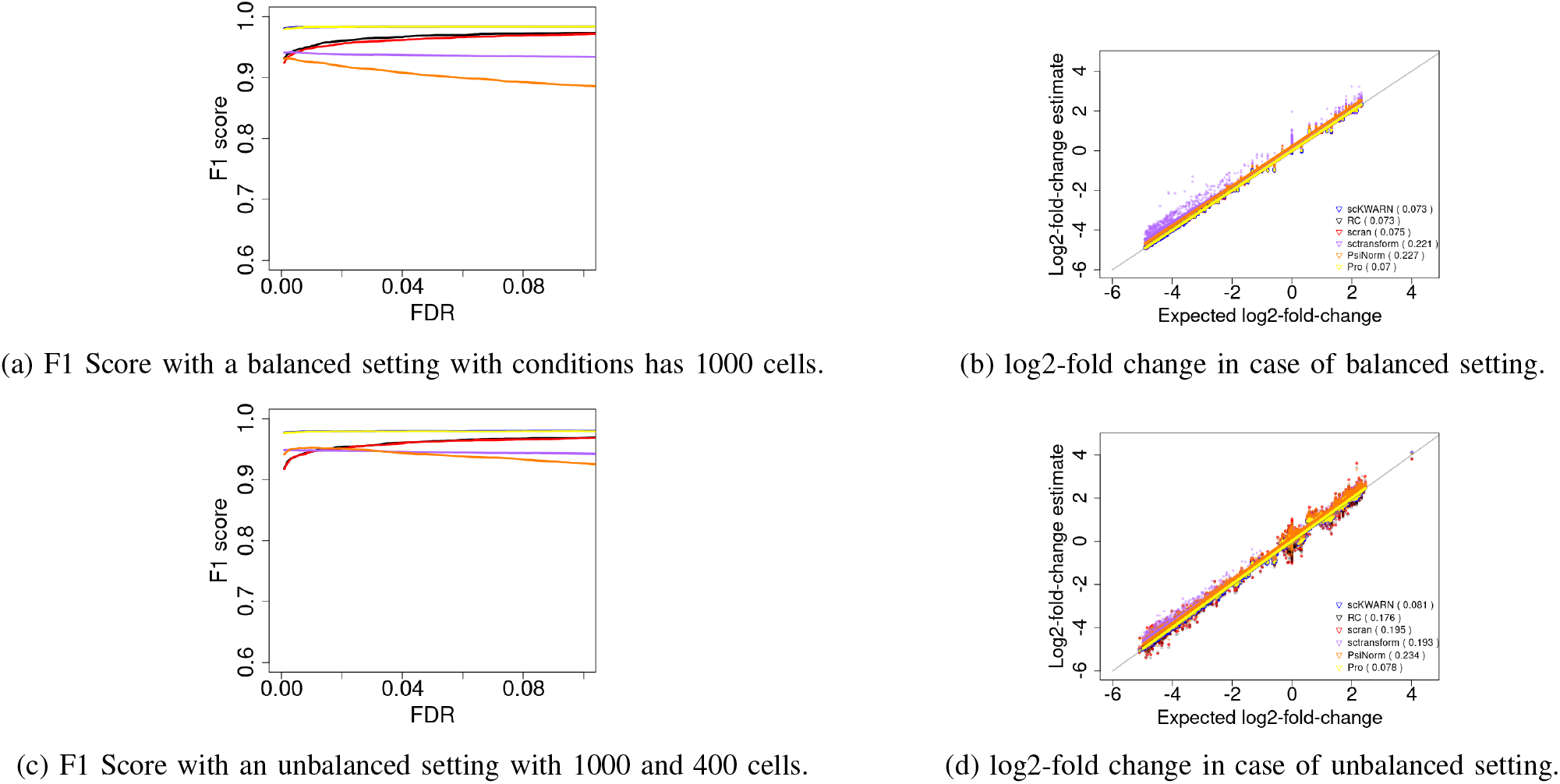
Performance analysis with state-of-the-art on PBMC33K dataset with Strong (30%) Differentially Expressed Genes.

#### 2) GSE29087

The second dataset used in this study was accessed from the Gene Expression Omnibus (GEO) database with accession number GSE29087. It contains 92 cells, comprising 48 mouse embryonic cells and 44 mouse embryonic fibroblast cells, with 22,928 genes. Genes with low expression counts across the libraries provide little information for differential expression (DE) analysis. These genes were filtered out if they were expressed in fewer than three cells. After removing the low-expression genes, 7,061 genes were included for DE analysis. Among these genes, 718, validated through qRT-PCR analysis, were taken as the gold standard DE genes, representing only a subset of true DE genes. In the current study, we used the top *G* genes ranked by each method as the gold standard genes. The performance was compared in three scenarios: none of cells, half of the cells, and all of cells downsized using a uniform random distribution0020with a minimum of 0.2 and a maximum of 1. When no cells were downsized and when half of the cells were randomly downsized, sctransform performed best in detecting DE genes, followed by the proposed approach, as shown in Figure 4. scran had the lowest proportion, possibly due to the limited number of cells, which may not be sufficient for accurate estimation of the scale factor. When all cells were downsized, the proposed approach again achieved better detection of DE genes compared to the other methods, except for sctransform, as shown on the right side of Figure 4.

**Fig. 4:**
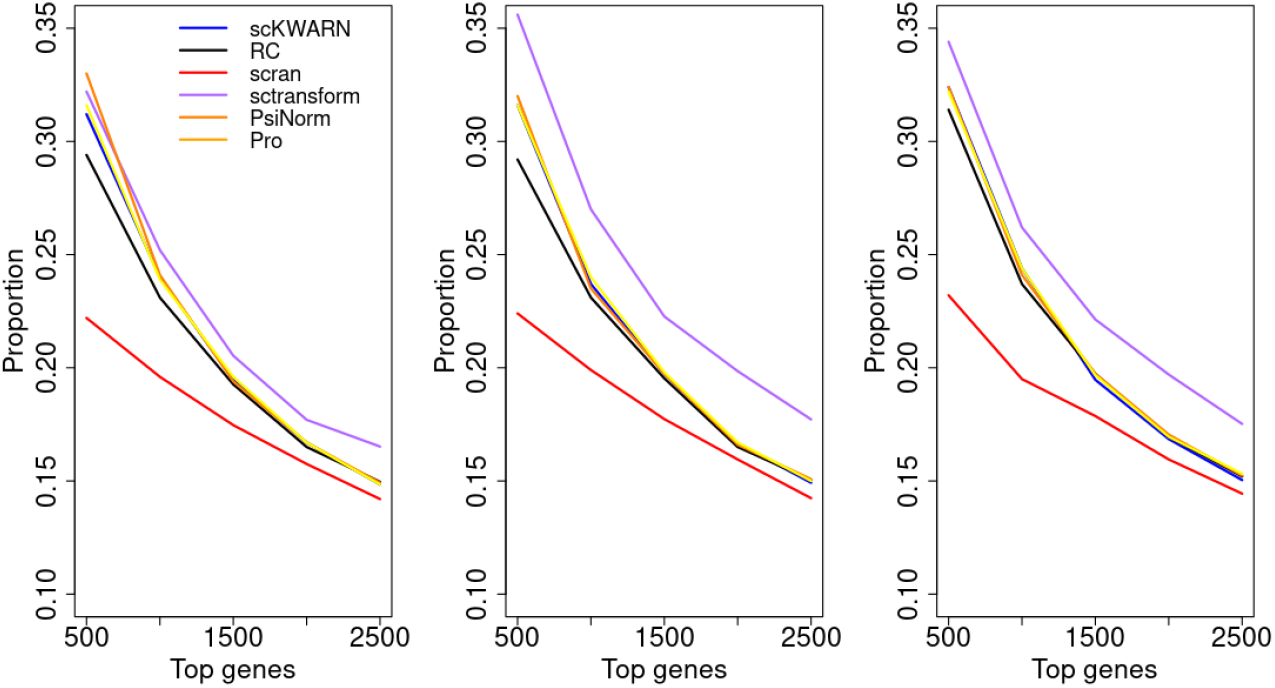
Performance analysis of proposed approach (Pro) with state-of-the-art on the [GSE29087](https://www.ncbi.nlm.nih.gov/geo/query/acc.cgi) dataset. Y axis represents the proportion of 718 gold standard genes in the top *G* genes on X axis. The performance was compared in three scenario: left plot with none of cells downsized,, middle plot with half of the cells downsized, and right plot with all cells downsized.

#### 3) GSE60361

The third dataset was accessed from GEO database with accession number GSE60361. It contains data from the somatosensory cortex (S1) and hippocampus CA1 area of juvenile (P22-P32) CD1 mice, comprising 33 males and 34 females. Cells were collected without selection, except for 116 cells obtained by FACS from 5HT3a-BACEGFP transgenic mice. A total of 76 Fluidigm C1 runs were performed, each attempting 96 cell captures and resulting in 3,005 highquality single-cell cDNAs, sequenced at an average depth of 14,000 reads per cell (with UMI) [22]. Similar to the first study, the present approach was tested on moderate DE and strong DE with balanced and unbalanced cell populations. For moderate DE, the percentage of genes varied from 5% to 15%; however, in strong DE, the percentage of genes varied from 15% to 30%. The balanced case involved each cell population having 1,000 cells, while the unbalanced case had one population with 1,000 cells and the other with 400 cells, similar to Study 1. In all cases, the present approach was compared with scKWARN, scran, RC, sctransform, and PsiNorm using performance metrics such as the F1 score and RMSE with true fold change. For moderate DE with balanced and unbalanced cases, as shown in Figure 5, the present approach achieved comparable or better performance in F1 scores and RMSE values. Figure 5 also shows that the present approach accurately estimates the true log-fold change while varying the percentage of DE genes within the cell population. For strong DE with balanced and unbalanced cases, as shown in Figure 6, the performance of the present approach was comparable to scKWARN and outperformed the rest of the normalization methods in F1 scores and RMSE values.

**Fig. 5:**
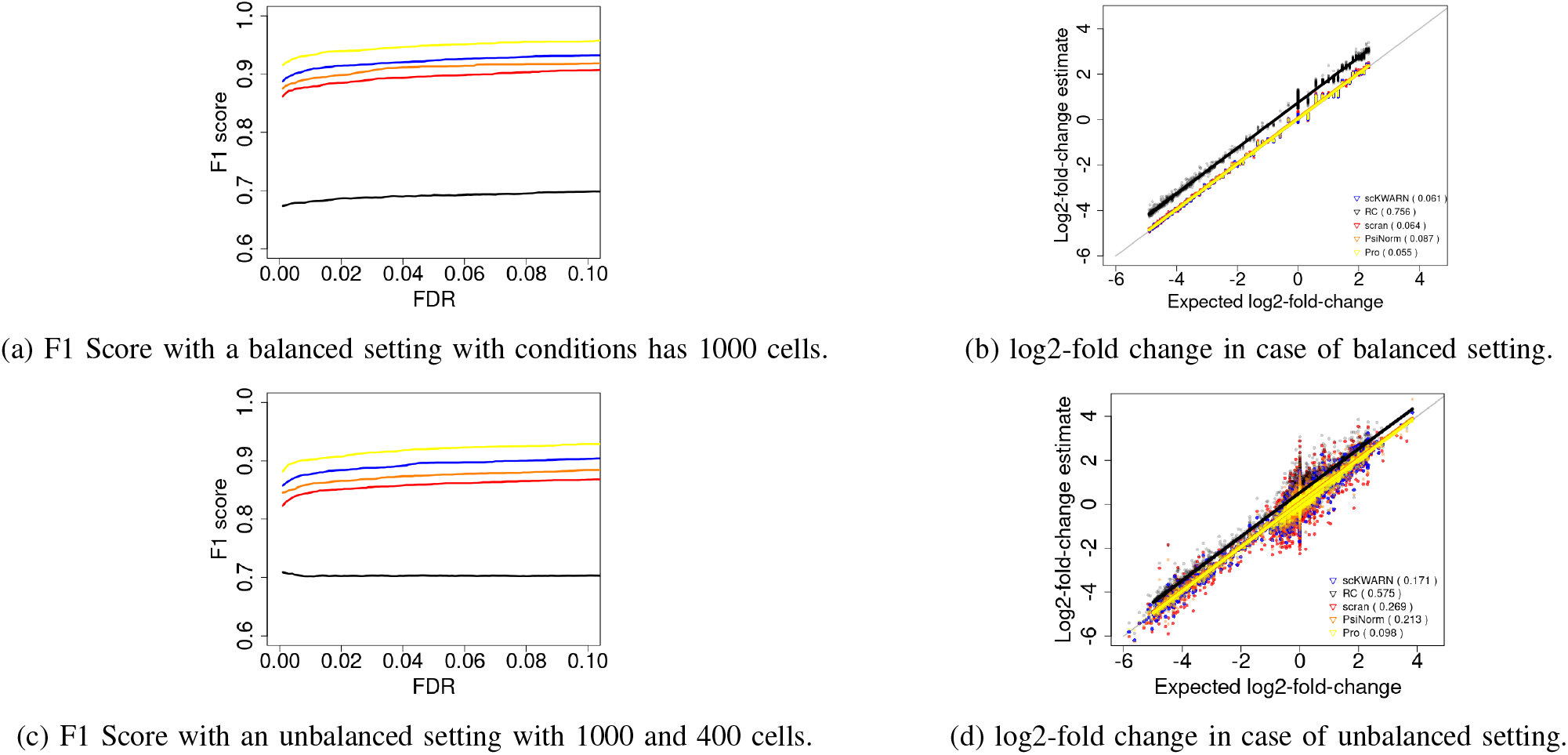
Performance analysis with state-of-the-art on GSE60361 dataset with Moderate (10%) Differentially Expressed Genes.

**Fig. 6:**
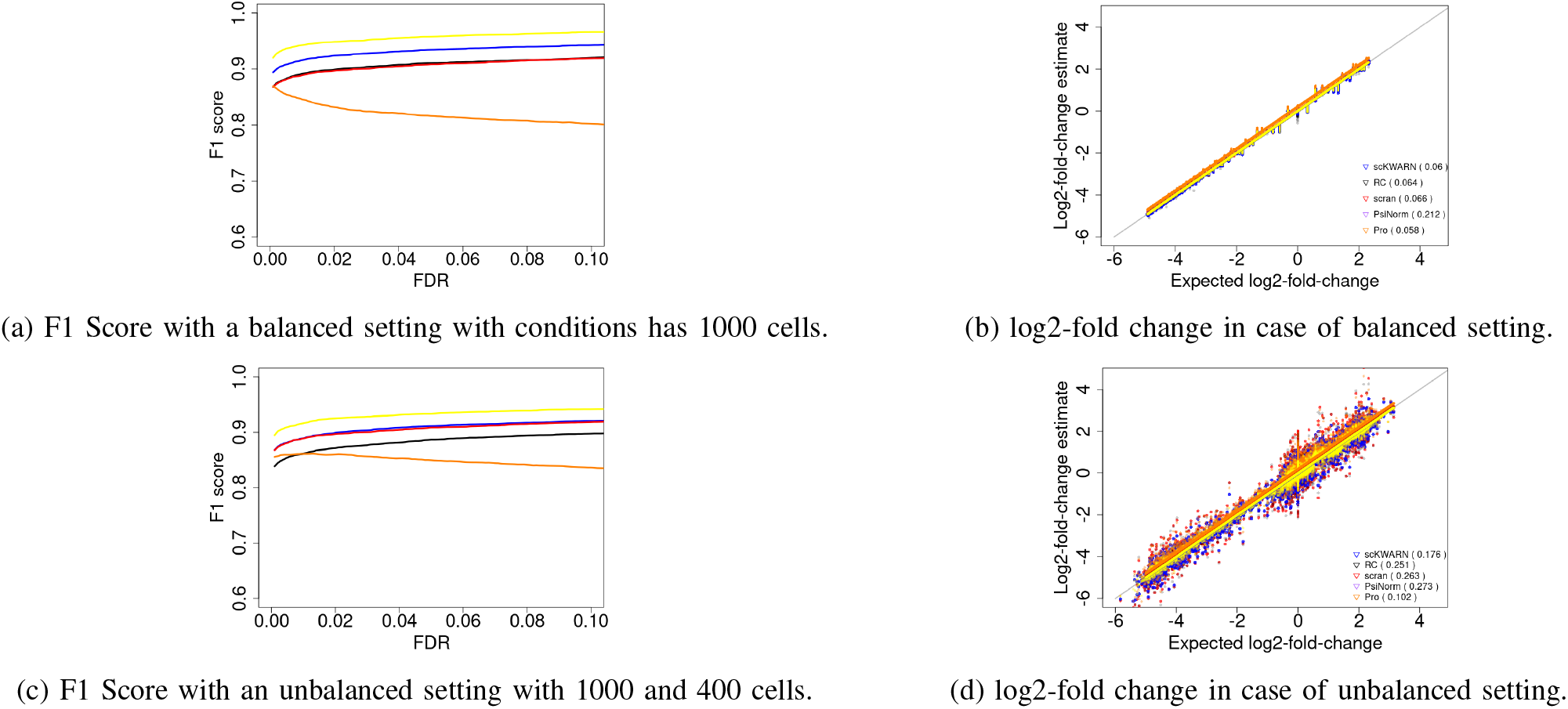
Performance analysis with state-of-the-art on GSE60361 dataset with Strong (30%) Differentially Expressed Genes.

### B. Simulated Datasets

The simulated data were generated using a negative binomial distribution under two cases: a linear relationship between the mean and count-depth [1], and no linear relationship between the mean and count-depth [8]. One hundred simulated datasets were generated for each case, consisting of 3,000 genes across three distinct cell groups. The effects of noise, such as library size, dropout rate, and RNA composition, were studied in both balanced and unbalanced settings. The simulated data were also analyzed by varying the percentage of DE genes, ranging from moderate to strong DE.

#### 1) Simulation I (SIM-I)

The first simulation setting, *y*_*gj*_, is generated from a negative binomial distribution with dispersion 0.1 and mean:

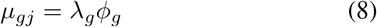

Where

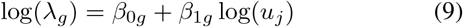

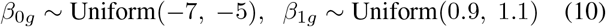

The parameter, *ϕ*_*g*_ = 1 is chosen for all genes excluding the DE genes in each of the three distinct cell groups. For DE genes, *ϕ*_*g*_ = 8*p* is chosen, where *p* follow the Bernoulli distribution:

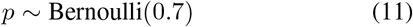

For altering library sizes of cells, we considered:

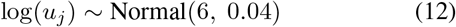

In the case of RNA compositions, 5% genes are randomly chosen (excluding DE genes) and assumed to be highly expressed genes by multiplying their expression levels with a positive constant chosen from the set *{*12, 13, …, 20*}*. These 5% highly expressed genes dominate 20% to 60% of library sizes. To model dropout rates, we increased the dropout with an additional probability of 10% to 30% as follows:

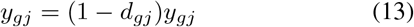

Where

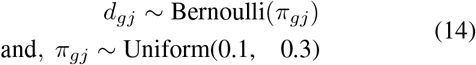

#### 2) Simulation II (SIM-II)

In the second simulation setting, *y*_*gj*_, is also generated from negative binomial distribution with mean *µ*_*gj*_ as of (8) with similar dispersion value of 0.1. The parameter *λ*_*g*_ follow a gamma distribution, specifically:

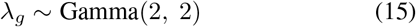

The parameter, *ϕ*_*g*_ is similar to that in condition SIM I. In the case of altering library sizes of cells, the mean of the negative binomial distributions were set as:

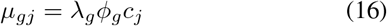

Where

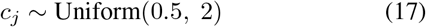

In the case of RNA compositions, 5% genes are randomly chosen (excluding DE genes) similar to SIM-I and assumed as highly expressed genes by multiplying a positive constant chosen from the set *{*12, 13, …, 20*}* with probability of 1*/*9. The 5% highly expressed genes dominate 20% to 60% of library sizes. For dropout rates, we also increased dropout with an additional probability 10% to 30% as of (13) in SIM-I.

The performance on the simulated datasets was assessed using four different metrics: F1 score, sensitivity, specificity, and bias. The R package [MAST](https://www.bioconductor.org/packages/release/bioc/html/MAST.html) was used to identify DE genes, and the difference between the true and estimated log 2 fold change was measured in terms of bias. The F1 score provides a balance between precision and recall. Additionally, sensitivity and specificity were used to assess the performance of the proposed approach. The performance was evaluated in terms of varying library size, RNA composition, and dropout for SIM-I and SIM-II. Figure 7 shows the performance of the present approach compared with state-of-the-art methods by altering the library size. As shown in Figure 7, the present approach exhibited less bias in both SIM-I and SIM-II, with improved overall performance. In the case of RNA composition, as shown in Figure 8, the present approach achieved a better F1 score, indicating a superior balance between precision and recall in both SIM-I and SIM-II. Finally, performance was evaluated based on dropout rate, as shown in Figure 9, for both SIM-I and SIM-II on moderately DE genes.

**Fig. 7:**
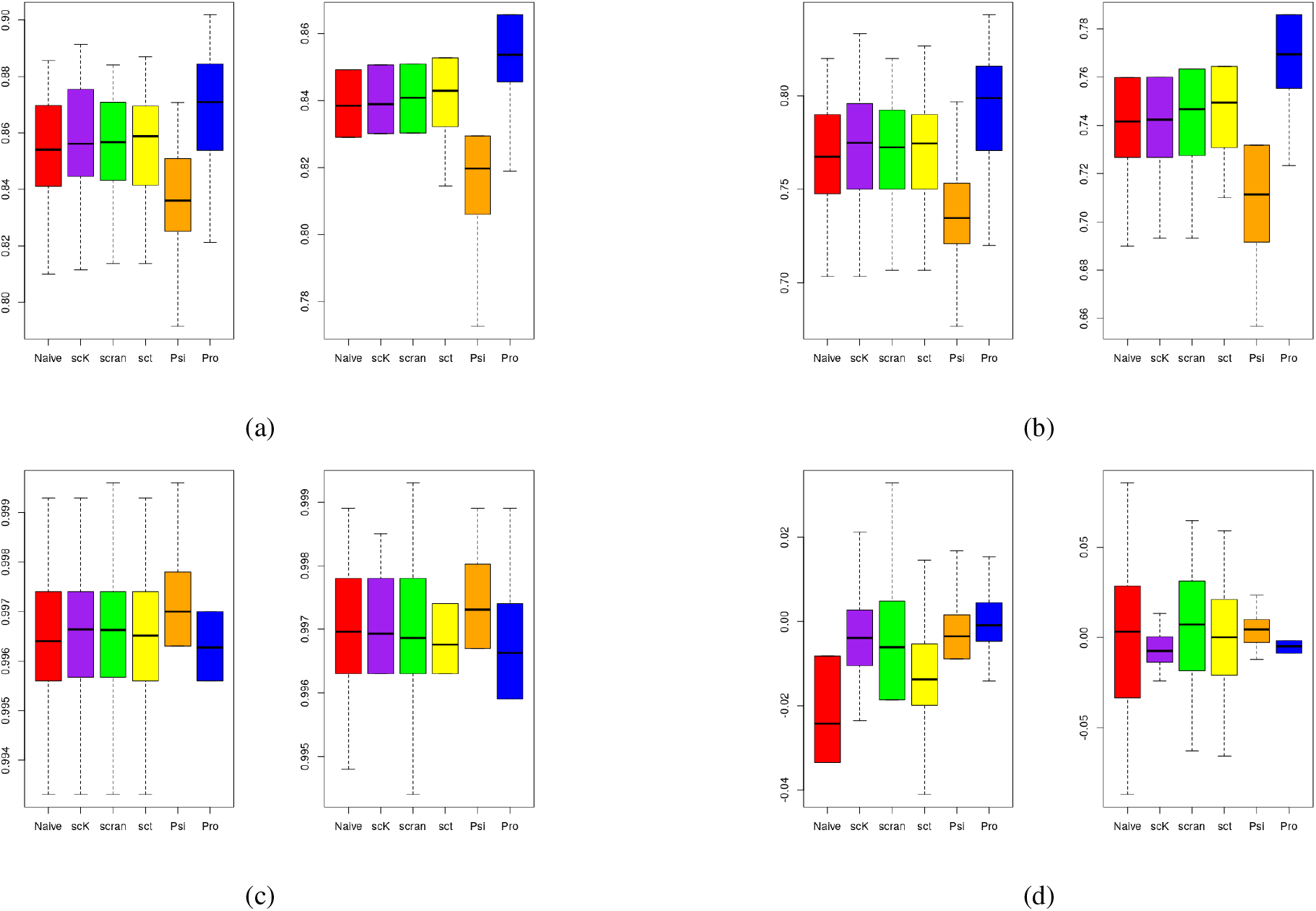
Performance analysis with state-of-the-art on the library size of dataset with Moderate Differentially Expressed Genes: (a) F1 score (left SIM-I and right SIM-II), (b) Sensitivity plot (left SIM-I and right SIM-II), (c) Specificity plot (left SIM-I and right SIM-II), and (d) Bias plot (left SIM-I and right SIM-II). Navie (RC), scK (scKWARN), sct(sctransform) and Psi(PsiNorm)

**Fig. 8:**
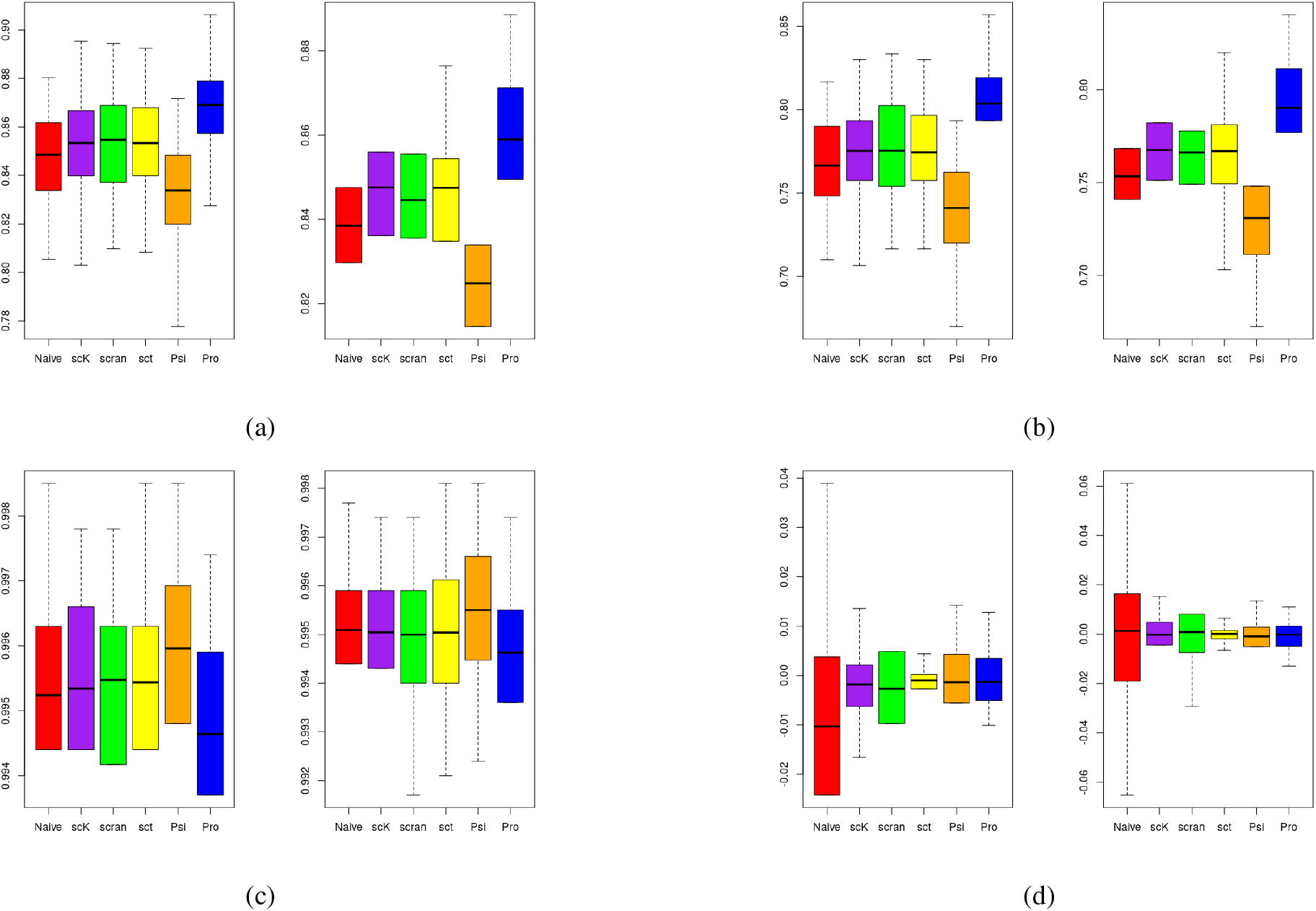
Performance analysis with state-of-the-art on the RAN composition with Moderate Differentially Expressed Genes: (a) F1 score (left SIM-I and right SIM-II), (b) Sensitivity plot (left SIM-I and right SIM-II), (c) Specificity plot (left SIM-I and right SIM-II), and (d) Bias plot (left SIM-I and right SIM-II).

**Fig. 9:**
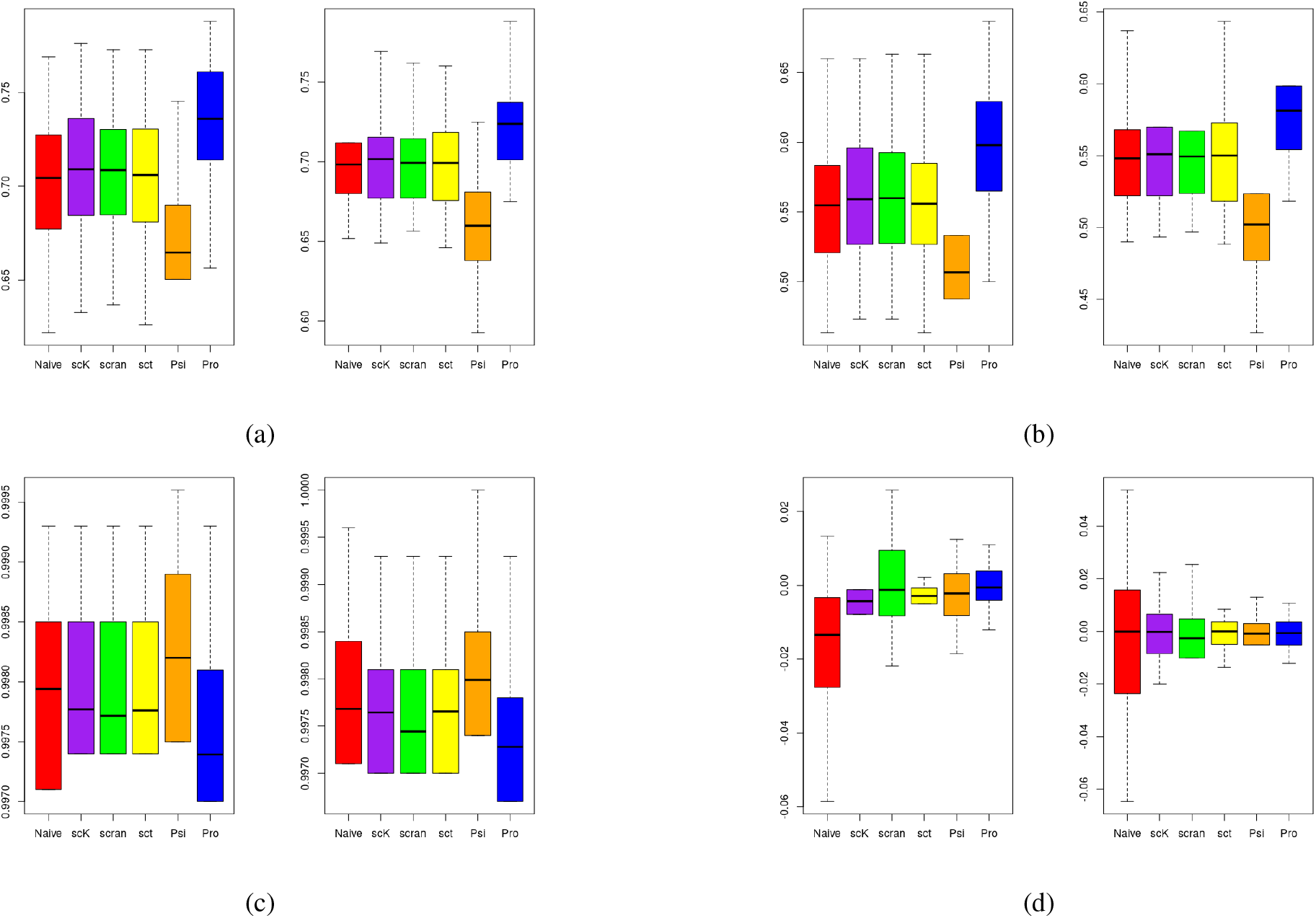
Performance comparison on the Dropout with Moderate Differentially Expressed Genes: (a) F1 score (left SIM I and right SIM II), (b) Sensitivity plot (left SIM-I and right SIM-II), (c) Specificity plot (left SIM-I and right SIM-II), and (d) Bias plot (left SIM-I and right SIM-II).

The comprehensive study of both real and simulated data shows that scran and PsiNorm have high RMSE when cell populations were unbalanced in both moderate and strong DE levels. The proposed approach and scKWARN consistently achieved low RMSE values across all three settings when biases were introduced by altering the library size, RNA composition, and dropout rates. sctransform estimates genespecific scaling factors, and its performance highly depends on the fitting of the model and whether the simulation settings align with their underlying assumptions.

## IV. Computational Performance

The computational performance of the present approach is presented in Table I. The execution time of the present approach is comparable to state-of-the-art methods. sctransform has a longer computation time because it estimates the genespecific scale factor by explicitly fitting each gene with a generalized linear model.

**TABLE 1:** Comparison of Runtime (In Seconds)

| Datasets | RC [1] | scran [8] | scKWARN [13] | scransform [1] | PsiNorm [18] | Proposed |
| --- | --- | --- | --- | --- | --- | --- |
| PBMC33K | 1.10 | 22.43 | 57.68 | 301.45 | 3.12 | 75.23 |
| GSE29087 | 0.03 | 0.41 | 1.67 | 38.32 | 0.17 | 7.14 |
| GSE60361 | 0.03 | 0.44 | 1.67 | 40.32 | 0.19 | 9.14 |
| SIM-I and SIM-II | 0.07 | 1.91 | 3.53 | 35.21 | 0.09 | 5.03 |

## V. Conclusion

This paper presents a robust normalization method for scRNA-seq data using PLS regression. Variability between conditions was minimized by multiplying with the average correlation. Biases in library size were reduced by modeling the scale factor with adaptive fuzzy weights using the upper and lower quintiles of the data. The effectiveness of the proposed approach was validated using real and simulated datasets and was compared with state-of-the-art methods across various performance metrics. The results show that the proposed approach effectively corrects biases due to library size, RNA composition, and dropout in scRNA-seq datasets.

## Notes

### Competing Interest Statement

The authors have declared no competing interest.

